# Using rapid prototyping to choose a bioinformatics workflow management system

**DOI:** 10.1101/2020.08.04.236208

**Authors:** Michael J. Jackson, Edward Wallace, Kostas Kavoussanakis

**Affiliations:** EPCC, The University of Edinburgh, United Kingdom; School of Biological Sciences, The University of Edinburgh, United Kingdom

**Author notes:** Corresponding authors (MJ), (EW).

## Abstract

Workflow management systems represent, manage, and execute multi-step computational analyses and offer many benefits to bioinformaticians. They provide a common language for describing analysis workflows, contributing to reproducibility and to building libraries of reusable components. They can support both incremental build and re-entrancy – the ability to selectively re-execute parts of a workflow in the presence of additional inputs or changes in configuration and to resume execution from where a workflow previously stopped. Many workflow management systems enhance portability by supporting the use of containers, high-performance computing systems and clouds. Most importantly, workflow management systems allow bioinformaticians to delegate *how* their workflows are run to the workflow management system and its developers. This frees the bioinformaticians to focus on the content of these workflows, their data analyses, and their science.

RiboViz is a package to extract biological insight from ribosome profiling data to help advance understanding of protein synthesis. At the heart of RiboViz is an analysis workflow, implemented in a Python script. To conform to best practices for scientific computing which recommend the use of build tools to automate workflows and to re-use code instead of rewriting it, the authors reimplemented this workflow within a workflow management system. To select a workflow management system, a rapid survey of available systems was undertaken, and candidates were shortlisted: Snakemake, cwltool and Toil (implementations of the Common Workflow Language) and Nextflow. An evaluation of each candidate, via rapid prototyping of a subset of the RiboViz workflow, was performed and Nextflow was chosen. The selection process took 10 person-days, a small cost for the assurance that Nextflow best satisfied the authors’ requirements. This use of rapid prototyping can offer a low-cost way of making a more informed selection of software to use within projects, rather than relying solely upon reviews and recommendations by others.

**Author summary:** Data analysis involves many steps, as data are wrangled, processed, and analysed using a succession of unrelated software packages. Running all the right steps, in the right order, with the right outputs in the right places is a major source of frustration. Workflow management systems require that each data analysis step be “wrapped” in a structured way, describing its inputs, parameters, and outputs. By writing these wrappers the scientist can focus on the meaning of each step, which is the interesting part. The system uses these wrappers to decide what steps to run and how to run these, and takes charge of running the steps, including reporting on errors. This makes it much easier to repeatedly run the analysis and to run it transparently upon different computers. To select a workflow management system, we surveyed available tools and selected three for “rapid prototype” implementations to evaluate their suitability for our project. We advocate this rapid prototyping as a low-cost (both time and effort) way of making an informed selection of a system for use within a project. We conclude that many similar multi-step data analysis workflows can be rewritten in a workflow management system.

## Introduction

Bioinformatics data analysis takes many steps, and a crucial but frustrating part of bioinformatics work is to run the right processing steps, in the right order, on the right data, reliably [1]. Usually these steps will involve disparate pieces of software from different sources, all run from the command line. For example, high-throughput sequencing data analysis may involve demultiplexing, trimming, cleaning, alignment, de-duplication, base quality score recalibration, and quantification. Phylogenetic analysis may involve selecting sequences, multiple sequence alignment, alignment trimming, and tree inference. Image analysis can also involve many steps applied to large numbers of images. Success in these multi-step data analyses generally requires writing a script to automate the steps. However, traditional shell scripts and even Makefiles have limited error reporting, are hard to debug, can be hard to restart after they go wrong, and can be challenging to move from one computer architecture to another. For example, bash scripts do not support re-entrancy or incremental build unless these functionalities are explicitly implemented by their authors, which can be a non-trivial development activity.

Workflow management systems – systems to represent, manage and execute analyses – address these problems [2-4]. They can provide a common language for describing analysis workflows, contributing to reproducibility and the building of libraries of reusable components. They can support both incremental build and re-entrancy, providing the ability to selectively re-execute parts of a workflow in the presence of additional inputs or changes in configuration and the ability to resume execution from where a workflow previously stopped. Many workflow management systems provide support to exploit software containers and package managers, high-performance computing systems and clouds. Most importantly, workflow management systems allow bioinformaticians to delegate how their workflows are run to the workflow management system, and its developers, freeing the bioinformaticians to focus on their science.

In this article, we describe the process that we used for selecting a workflow management system for our ribosome profiling software, RiboViz [5]. While Leipzig [4] offers advice on choosing a workflow management system based on the qualities of classes of workflow management system, we used an approach to selection focused on both the popularity of the candidate tools within the bioinformatics community and on the specific merits of the candidate tools in the context of our project’s specific requirements.

To select a workflow management system, a rapid survey of available workflow management systems was undertaken and candidates were shortlisted: Snakemake (https://github.com/snakemake/snakemake) [6], cwltool (https://github.com/common-workflow-language/cwltool), and Toil (https://github.com/DataBiosphere/toil) [7] (chosen for their implementation of the Common Workflow Language [8], a language for describing workflows) and Nextflow (https://github.com/nextflow-io/nextflow) [9]. An evaluation of each candidate, via rapid prototyping of a subset of the RiboViz workflow, was performed and Nextflow was chosen. Our approach is summarised in Table 1

**Table 1.** Rapid prototyping to select software for a project.

| Step | Effort (person-days) | Tasks |
| --- | --- | --- |
| Conduct a rapid survey to identify potential software | 1-2 | Discover what software is available which may suit your requirements, which is in common use and is well-regarded within the community. |
| Shortlist candidates for rapid prototyping | 1 | Draw up a shortlist of candidates based on their popularity, whether they have an acceptable software licence, and whether |
|  |  | they are well-established, stable, and with evidence of a future. |
| Rapidly prototype a subset of functionality within each candidate | 2-3 (per candidate) | Rapidly prototype a subset of functionality required by the project and assess each candidate's suitability against objective project-specific criteria (required and functionality, supported platforms, etc.) and general subjective criteria (ease of install, ease of implementation, quality of supporting documentation). |

Our evaluation used rapid prototyping for three reasons. Firstly, using the candidate systems, and their documentation, would provide more insight into their ease of use, their capabilities, and the quality of their supporting documentation than could be ascertained by solely reading their documentation. Secondly, focusing on implementing our workflow, would give us more insight into these qualities than solely working through tutorial examples specifically designed by the developers of the systems to demonstrate their software. And, thirdly, whatever system we adopted, we would have the corresponding prototype to build upon.

Though our focus was on selecting a workflow management system, the use of rapid prototyping offers a low-cost way of making a more informed selection of software to use within projects, rather than relying solely upon reviews and recommendations by others.

The intent of this article is not to make a recommendation as to the use of a specific workflow management system for all bioinformatics projects. Nor is this article intended to claim that using rapid prototyping is suitable for the selection of all software or for all projects. Rather, it is to demonstrate how we used rapid prototyping to select a workflow management system that met the specific requirements of our project, and to discuss our experiences with the workflow management systems that we considered.

### RiboViz and the requirement for a workflow management system

RiboViz is a high-throughput sequencing analysis pipeline specialised for ribosome profiling data. RiboViz takes raw data from sequencing machines; estimates how much each part of RNA is translated into protein and how the amount of translation is controlled by the code of that RNA; and produces analysis data, tables, and graphs.

RiboViz is under active development by The Wallace Lab and EPCC at University of Edinburgh, The Shah Lab at Rutgers University and The Lareau Lab at University of California, Berkeley. The source code is available on GitHub (https://github.com/riboviz/riboviz), under an Apache 2.0 open source licence.

At the heart of RiboViz is an analysis workflow to process ribosome profiling data across several samples, whose information along with all parameters for processing is described in a single input YAML file. Sample-specific read data can be provided as separate (fastq) input files or within a multiplexed input file. This workflow invokes a series of steps per sample (for example, adapter trimming, rRNA and ORF alignment, trimming 5’ mismatches). In addition, there are some initial, sample-independent, steps (for example, creating rRNA and ORF indices). When all samples have been processed, results from the sample-specific analyses are aggregated and summarised. Some steps (for example UMI extraction and deduplication) are conditional upon the nature of the samples and configuration parameters set by the user. The workflow does not include any loops. S1 Fig and S2 Fig show the steps of the RiboViz workflow that are invoked when processing demultiplexed samples and when processing multiplexed samples, respectively.

Each discrete step in a sequencing analysis workflow corresponds to the invocation, via bash, of a command-line tool. Some of these tools are open source packages in widespread use within the bioinformatics community and include HISAT2 [10], Cutadapt [11], Samtools [12], Bedtools [13] and UMI-tools [14]. Other tools, implemented in Python and R (for demultiplexing multiplexed files, trimming reads and generating analysis data, tables, and graphs) have been developed by the RiboViz team.

The RiboViz analysis workflow was implemented in a Python script. Each time a command-line tool is invoked, a log file is created for each invocation, in which standard output and error is captured. A log file for the execution of the Python script itself is also created. Sample-specific data and log files are written to sample-specific directories. The Python script also logs all the commands executed via bash to a script which can be run standalone and which allows a specific analysis to be rerun outwith the Python script. The RiboViz Python script can be configured to run in a “dry run” mode whereby it will validate its configuration, check that input files exist and output this complete bash script without executing the steps.

The design of the RiboViz Python script follows the majority of van Vliet’s seven quick tips for analysis scripts [15], tips which are generally applicable beyond neuroimaging. The development of RiboViz has also been strongly guided by Wilson et al.’s best practices for scientific computing [16]. We use version control, unit test libraries, and, to help manage collaboration, an issue tracker. We turn bugs into test cases and endeavour to write programs for people. Their recommendation to “Write code in the highest-level language possible” motivated our migration to Python from a previous implementation of the analysis as a bash script.

However, as our Python script evolved, we were aware that we were adding more features related to managing the invocation of the analysis steps, rather than the nature of these steps themselves – we were implementing a custom workflow management system for RiboViz. This was problematic for several reasons. Our code was becoming more difficult to maintain as it evolved to accommodate additional requirements which were not envisaged when its implementation began in 2016. Our code did not support re-entrancy or incremental build, both of which we viewed as essential for implementing workflows to process large datasets. Nor did our code support parallel execution of the workflow which would be necessary to support the future execution on RiboViz on large-scale datasets. Implementing these would have incurred significant development effort, effort which would be better spent implementing the steps within the workflow, the science itself.

It was time for us to adopt two more of the Wilson et al. best practices, to “Use a build tool to automate workflows” and to “Re-use code instead of rewriting it”, that is, to use an off-the-shelf workflow management system, the adoption of which, we estimated, would incur significantly less effort than implementing re-entrancy, incremental build and support for parallel processing ourselves.

### A survey of available workflow management systems to shortlist candidates

We first conducted a rapid survey of available workflow management systems to shortlist candidates for rapid prototyping. The criteria we used to select candidates for shortlisting are summarised in Table 2.

**Table 2.** Shortlisting criteria.

| Criteria | Description |
| --- | --- |
| Popularity | The system seems to be in common use and is well-regarded within the bioinformatics community. The system is likely to be practically usable. |
| Free and open source licence | The system is free and has an open source licence, as RiboViz itself is free and open source. |
| Well-established, stable and with a future | The system has been around for at least a year, has regular releases and evidence that it is actively maintained, developed, and supported. Development of the system is unlikely to stop after we migrate to it. |

We started by conducting web searches to find out what systems, and existing surveys of systems, were available using combinations of the terms “workflow management system” and “bioinformatics,” “survey” and “list”. In keeping with our pragmatic, low-cost, approach, a systematic literature review was not undertaken as our goal was not to produce a comprehensive survey of every workflow management system available, but to, in a rapid way, identify which systems are in common use, and are well-regarded, within the bioinformatics community.

We consulted existing surveys of workflow management systems. Leipzig [4] provides a comprehensive review of the types of workflow management systems available and the pros and cons of each type, with examples of 16 workflow management systems. Di Tommaso et al. [17] lists 106 pipeline frameworks and libraries and 30 workflow platforms. The Common Workflow Language wiki [18] has a stated-incomplete list of 279 computational data analysis workflow systems. To avoid being rendered indecisive through over-choice, we proceeded to look for more subjective recommendations and discussions of the pros and cons of specific systems.

Baichoo et al. [19] describe a case study of a project, H3ABioNet, who sought a workflow management system both in common use within the bioinformatics community and consistent with the skill-set of their project and their collaborators and which were both popular and portable. Their project selected both Common Workflow Language and Nextflow as these were deemed to meet the project’s criteria. CWL is a language for describing workflows and there are a number of workflow management systems that can execute workflow written in CWL (Baichoo et al. authors used its reference implementation, cwltool (https://github.com/common-workflow-language/cwltool)).

An online discussion on “Given the experience of others writing bioinformatic pipelines, what are the pros/cons of Toil vs Snakemake vs Nextflow?” [20] provides a wealth of opinion on workflow management systems. Nextflow was viewed very positively, for its ease of use, reporting, portability, documentation, community, and error reporting. However, its reliance on Groovy (https://groovy-lang.org/), a Python-style scripting language that can be run on the Java platform, was perceived to be daunting and Nextflow’s error messages were deemed to be quite cryptic. Snakemake was viewed positively for its ease of use, the concision of its workflows, and for being based on Python. However, its documentation was felt to be lacking and it was not considered to be as flexible as Nextflow. The use of Workflow Definition Language [21] and CWL, in conjunction with workflow management systems that implement these (for example Cromwell (https://github.com/broadinstitute/cromwell/) [22]), were viewed positively for allowing the implementation of simple, portable, workflows and, in the case of WDL, for supporting subworkflows. The use of such languages was deemed challenging for more complex workflows, or those that require custom code to perform specific tasks.

In a poll on workflow management systems, tools, platforms, languages and specifications [23], completed by 36 respondents, 44.4% voted for Nextflow, 22.4% for SevenBridges (https://www.sevenbridges.com/), and 19.4% for CWL.

From the available options, we decided to choose the two systems that had, in our impression, been most frequently, and positively, mentioned within the foregoing papers and the online discussion, had received the most votes in the poll, and had been recommended by our team’s bioinformaticians and their colleagues. We only considered free and open source systems as RiboViz itself is free and open source (this precluded SevenBridges, a commercial system). Our shortlisted candidates were Snakemake and Nextflow. CWL had also been frequently and positively mentioned so we chose both its reference implementation, cwltool, and one of its production implementations, the aforementioned Toil. We chose Toil over Cromwell as Cromwell was listed as a “partial”, not “production”, implementation on the CWL web site.

It was also important that we adopt a workflow management system that was well-established, stable and with a future [24]. We did not want to migrate to a system only for development around that system to stop. To assess the stability of and development activity around each tool, we reviewed statistics from their open source repositories and the number of web search results for them (Table 3).

**Table 3.** Statistics on the shortlisted workflow management systems.

| Workflow management system | Software licence | Project start date | Last updated | Number of contributors | Search results |
| --- | --- | --- | --- | --- | --- |
| Snakemake | MIT | 2013 | Week beginning 24/04/2020 | 122 | 31,100 |
| Nextflow | Apache 2.0 | 2013 | Week beginning 24/04/2020 | 230 | 31,000 |
| cwltool | Apache 2.0 | 2014 | 28/02/2020 | 72 | 2,440 |
| Toil | Apache 2.0 | 2011 | 28/02/2020 | 81 | 63,600 |

“Last updated” and “Number of contributors” were documented on 28 February 2020. “Search results” are the number of search results for the search term “<workflow management system> bioinformatics” on Google on 21 July 2020.

All shortlisted systems have permissive open source licenses, have been in existence for several years, have many contributors and are regularly updated. Overall, this survey gave us confidence that the candidate systems were being widely and actively used, developed, and supported, and will continue to be so for the foreseeable future.

### Evaluation of candidates via rapid prototyping

Once a shortlist had been drawn up, we carried out an evaluation of each candidate system via rapid prototyping. This allowed for a more detailed evaluation as to whether each of the candidate systems met our requirements as well as to assess how easy it is to use the systems and the perceived quality and utility of their supporting documentation.

Our evaluation focused on rapidly prototyping a subset of the RiboViz workflow into each system. There were three reasons for this. Firstly, using each system, and their documentation, would provide more insight into their ease of use, their capabilities, and the quality of their supporting documentation than could be ascertained by solely reading their documentation. Secondly, focusing on implementing our workflow would give us more insight into these qualities than solely working through tutorial examples specifically designed by the developers of the systems to demonstrate their software. And, thirdly, whatever system we adopted, we would have the corresponding prototype to build upon.

2-3 person-days were allotted to each system. If nothing productive could be implemented within that period, then the system would be left and the next considered.

Our evaluation criteria are shown in Table 4.

**Table 4.**
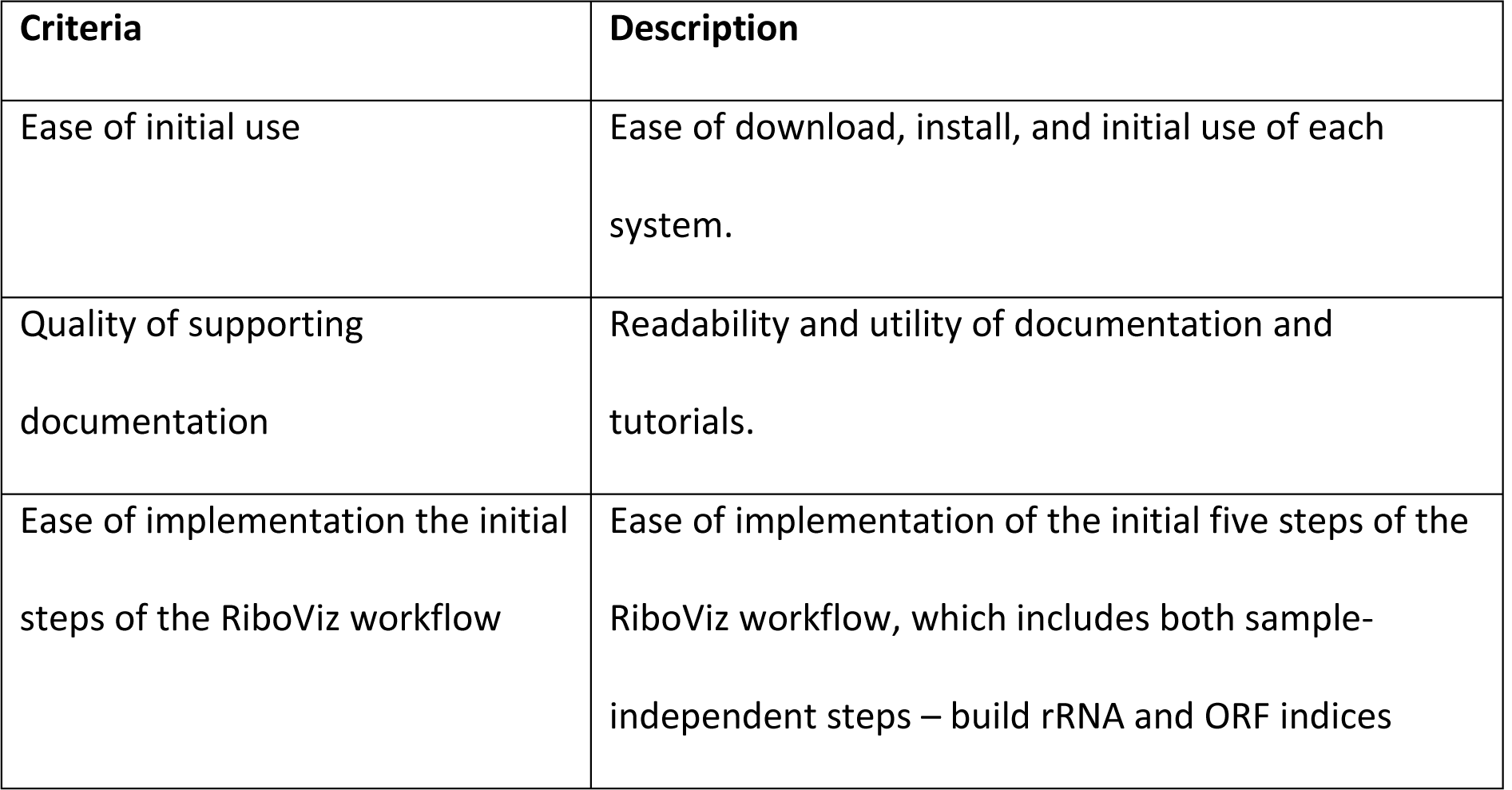
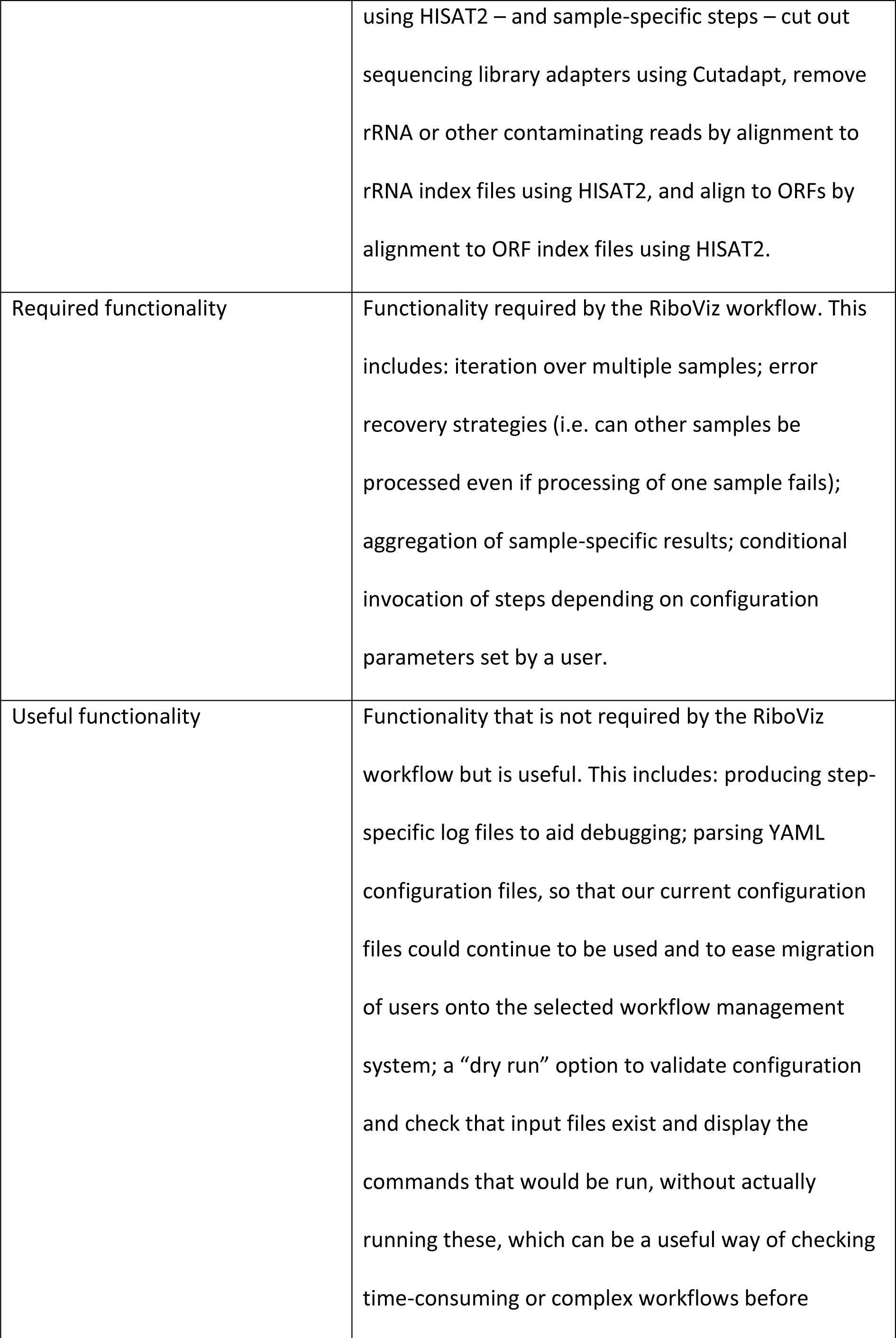
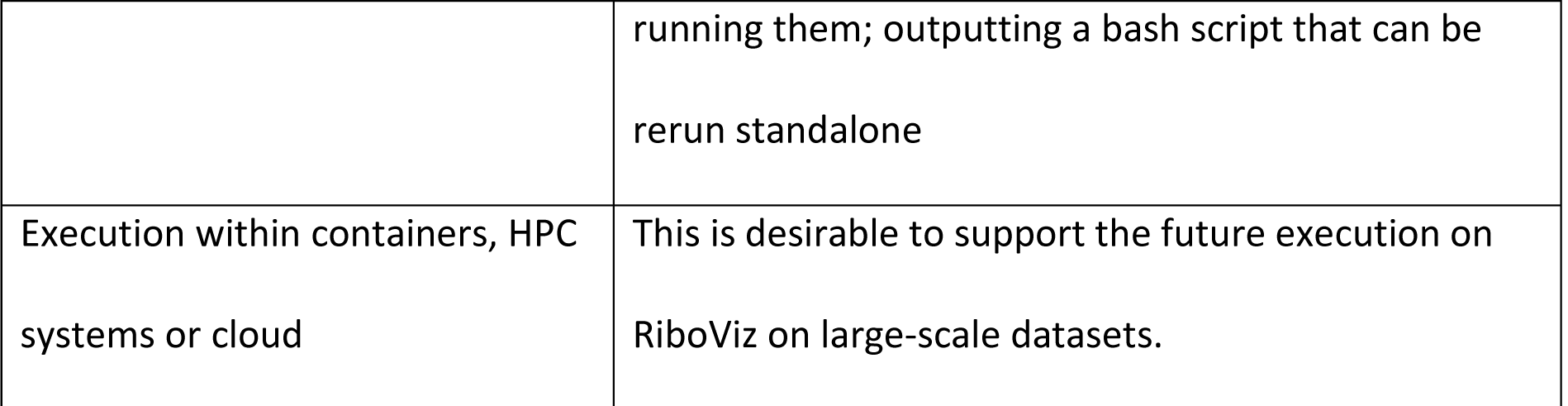
Evaluation criteria.

It will be noted that the first three criteria are subjective, and necessarily so [25]. Ease of use, readability of documentation and ease of implementation are very much dependent upon the skills, knowledge, and experience of those who will use a system and its supporting resources. For RiboViz, users are expected to be familiar with bash command-line tools and developers familiar with development of bash, Python and R scripts under Linux. We sought a system that would enable us, and our user community, to implement, maintain and extend our workflow in a way that is easier than at present. Table 5 summarises how each tool met our evaluation criteria.

**Table 5.** Summary of workflow management systems and objective evaluation criteria.

| Criteria | Snakemake | cwtool | Toil | Nextflow |
| --- | --- | --- | --- | --- |
| Installer | conda | Python pip | Python pip | conda |
| Number of RiboViz processing steps implemented | 14 | 3 | 3 | 5 |
| Time (person-days) to implement steps | 1 | 1<sup>1</sup> | 1<sup>1</sup> | 2 |
| Iteration over multiple samples | Yes | See note<sup>2</sup> | See note<sup>2</sup> | Yes |
| Error recovery strategies | Yes (“keep going” parameter) | See note <sup>2</sup> | See note <sup>2</sup> | Yes (step-specific error strategies) |
| Aggregation of sample-specific results | Yes | See note <sup>2</sup> | See note <sup>2</sup> | Yes |
| Conditional execution of steps | Yes (step-specific Python conditions) | No | No <sup>3</sup> | Yes (step-specific “when” clauses) |
| Step-specific log files | Yes (must be captured explicitly by each step) | See note <sup>2</sup> | See note <sup>2</sup> | Yes (automatically captured) |
| YAML configuration files | Yes | See note <sup>2</sup> | See note <sup>2</sup> | Yes |
| “dry run” option | Yes | See note <sup>2</sup> | See note <sup>2</sup> | No (but configuration can be implemented) |
| Output a rerunnable bash script | No (outputs a summary file with bash commands) | See note <sup>2</sup> | See note <sup>2</sup> | No (outputs step-specific bash scripts) |
| Execution within containers, HPC systems and cloud | Yes | Containers only | Yes | Yes |

Summary of how each workflow management system met the RiboViz project’s objective evaluation criteria. ^1^ the time taken relates to writing CWL workflows, not cwltool or Toil-specific workflows. ^2^ these criteria were not explored as the decision had been made to not consider CWL further considering its lack of support for conditional execution of steps. ^3^ the lack of support for conditional execution is a restriction of CWL, not Toil.

#### Snakemake

Snakemake was easy to download and install, via the conda (https://docs.conda.io/en/latest/) package manager, and had a comprehensive tutorial. Snakemake adopts the same model of operation as the GNU Make (https://www.gnu.org/software/make/) automated build tool – users specify the output files they want to build, Snakemake looks for rules to create these output files and runs the commands (in Snakemake, bash commands or Python scripts) specified in these rules to create the output files. Rules can specify dependencies – files used by the commands to create the output files. If these files do not exist then Snakemake looks for rules to create these, and so on.

Snakemake is implemented in Python. Python code can also be embedded within a Snakefile, for example, to create file paths or validate configuration parameters.

Implementing steps from the RiboViz workflow was straightforward and a functional version of the complete RiboViz workflow (everything bar steps specifically to handle multiplexed files) was implemented in less than a person-day.

Snakemake provided all the required and useful functionality listed in our evaluation criteria. Snakemake provides a “keep going” configuration parameter which can be used to continue processing other samples if processing of one sample fails. Like Make, Snakemake supports incremental build and re-entrancy. Conditional behaviour can be executed via the use of Python conditions. Step-specific log files can be implemented, but Snakemake does not automatically capture these – the bash commands executed by each step explicitly need to redirect standard output and standard error streams into these log files. Like Make, Snakemake supports a “dry run” option that can check that input files exist and that displays the commands that would be run, without running these. As for Make, the ability to specify exactly the files to build can be useful for debugging.

While Snakemake does not output a bash script that can be run standalone, it can output a summary file with the commands submitted to bash for execution. This file could be parsed, and the commands extracted and constructed into a bash script.

Snakemake has support for running its jobs within containers, HPC systems and clouds.

#### Common Workflow Language, cwltool and Toil

Both cwltool and Toil were easy to install, via the Python pip (https://pip.pypa.io/) package manager. It was easy to run a CWL “hello world!” example via both. A comprehensive, step-by-step tutorial to the language is available (https://www.commonwl.org/user_guide) [26].

CWL tool wrappers, which describe the inputs and outputs of command-line tools, and job configuration files, which describe workflows, are written as YAML or JSON documents. JavaScript can be embedded for any additional computation that is required, for example to create file paths or validate configuration parameters.

Implementing three steps of the RiboViz workflow took a person-day. The “edit-run-debug” development cycle felt slow and painful, due to the richness of CWL and the occasionally cryptic error messages that arose during execution.

Conditional behaviour is not yet supported within CWL – a “Collecting use cases for workflow level conditionals” issue [27] was added in February 2020 to their 1.2 milestone, but, at time of writing (August 2020), this has no due date. The lack of conditional invocation means that CWL is not currently suitable for RiboViz, or for other projects that require input-dependent control of workflow structure. (A colleague had evaluated CWL about a year and a half ago and, while they felt that simple workflows showed promise, the lack of conditionals meant that they could not adopt CWL for their project. Similarly, we felt that CWL would not be suitable for RiboViz at this time.) This limitation could have been identified at the shortlisting stage, but we had to achieve a balance between how many criteria to consider during shortlisting and how many during our rapid prototyping. We (incorrectly as it turned out) assumed that support for conditional execution would be a fundamental feature of any workflow management system or languages, such as CWL, executed by them.

#### Nextflow

Nextflow was easy to download and install, via the conda package manager, and had a simple tutorial.

A Nextflow workflow has a structure analogous to a Makefile or Snakefile – it consists of a set of processes which define inputs, outputs and commands describing how to create the outputs from the inputs. However, Nextflow adopts a dataflow programming model whereby the processes are connected via their outputs and inputs to other processes, and processes run as soon as they receive an input. Unlike Snakemake, a user does not specify the files they want to create, rather, they declare their input files and related configuration and Nextflow continues to invoke processes until no process has any outstanding inputs. Nextflow is implemented in Java. Code written in Groovy, which runs under Java, can be embedded within a Nextflow workflow. This code can be used to, for example, create file paths or validate configuration parameters.

Implementing steps from the RiboViz workflow was straightforward and a functional version of the subset of the RiboViz workflow was implemented in two person-days. Nextflow’s documentation was comprehensive but additional concrete examples of how to implement common use cases, and an expanded tutorial, to implement a complex multi-step workflow, would have been helpful. We did, later, find a Nextflow 2017 workshop tutorial [28] but our understanding of Nextflow had, at that point, passed beyond its content. Despite this, writing a Nextflow workflow was easier than writing a CWL workflow.

Nextflow provided all the required and most of the useful functionality listed in our evaluation criteria. Each step can have an “error strategy” which indicates what to do if an error is encountered. This can be used to ensure that other samples are processed if processing of one sample fails, or to adjust process resource parameters if a reported error arose from a lack of memory or a time limit that was too low. Nextflow, like Snakemake, supports both incremental build and re-entrancy, via a “resume” option. Conditional execution of steps is supported via a “when” declaration. Unlike Snakemake it is not possible to specify the exact files to build, which can make debugging more challenging. However, every invocation of a step takes place in its own isolated subdirectory which includes a bash script with the command that was invoked, symbolic links to input files, output files, and files with the contents of the standard output and error streams. The step-specific bash scripts can be run within their step-specific directories which is useful for debugging the implementation of individual steps. These directories have auto-generated names but Nextflow allows the contents of these directories to be written into known locations with more readable names.

A “dry run”, analogous to that supported by Snakemake and Make, has been suggested in a Nextflow issue [29], but has not progressed due to challenges in implementing such a feature within a dataflow model. The Nextflow authors instead recommend using small datasets to validate scripts. It should be noted that it may be challenging to identify a small dataset that would allow adequate replication of the workflow’s behaviour in the presence of a full dataset.

Nextflow has support for running its jobs within containers, HPC systems and cloud.

### Selecting a workflow management system

We decided to adopt Nextflow for the following reasons. It was our subjective impression that Nextflow felt far richer than Snakemake both in terms of features and expressivity, and it was felt that these outweighed its lack of a dry-run feature. The execution of each step within isolated subdirectories is useful for debugging. While writing Nextflow workflows does require knowledge of Groovy, the authors, familiar with Python and R, did not find learning Groovy challenging. The fact that Nextflow was based on Java incurs no additional installation overhead for either users or developers compared to Snakemake – each can be installed using the conda package manager using a single command. Based on our impressions of their documentation, Nextflow’s built-in support for, and documentation around, containers, HPC systems and cloud, seemed more thorough than that of Snakemake (though we appreciate that this may change as both tools evolve).

### Completing the RiboViz workflow implementation in Nextflow

It took approximately five person-days to complete an implementation of the RiboViz workflow (including support for multiplexed files) within Nextflow. Our existing regression test framework for our Python script was used to validate the implementation of our Nextflow script.

The Nextflow implementation has been tested by the RiboViz development team on their own development platforms and also on EDDIE, The University of Edinburgh’s high performance computing cluster (https://www.ed.ac.uk/information-services/research-support/research-computing/ecdf/high-performance-computing).

Release 2.0 of RiboViz [30] includes the Nextflow implementation of the RiboViz workflow. The Python implementation of the RiboViz workflow will be deprecated in a future release. Nextflow has the nf-core collection of bioinformatics pipelines, a resource of open-source, reviewed, and validated Nextflow scripts implementing common data analyses [31]. The associated nf-core developer community (136 members as of 8 July 2020, https://nf-co.re/community) has some overlap with the Nextflow developers, but is primarily composed of bioinformaticians. Again, these provide strong evidence for a well-established system with a future and we will consider contributing RiboViz to nf-core in the future.

However, no choice of software should be permanently binding. Our positive experiences with Snakemake, and the small effort that would be required to complete the implementation of RiboViz into Snakemake, give us confidence that if we need to migrate from Nextflow to Snakemake in future, then this would be a relatively straightforward migration to undertake.

## Conclusion

In this article, we described the process that we devised for selecting a workflow management system for our ribosome profiling software, RiboViz. The use of rapid prototyping gave us a prototype from which to develop an implementation in our chosen workflow management system. Our approach took approximately 10 person-days for background reading, our survey and short-listing and rapid prototyping. In our view, this is a small cost for the assurance that our selected workflow management system meets our requirements, is well-established, widely and actively used, developed and supported, and will continue to be so for the foreseeable future. The use of Nextflow provides us with an implementation of RiboViz that is more maintainable, more portable, and which will allow us to exploit the power of distributed computing resources in the analysis of large-scale datasets.

Reiter et al. [32] also recommend rapid prototyping, in this case for migrating to a chosen workflow management system. In particular they recommend that “When building a workflow for the first time, creating an initial workflow based on a subset of your sample data can help verify that the workflow, tools, and command line syntax function at a basic level. This functioning example code then provides a reliable workflow framework free of syntax errors which you can customize for your data without the overhead of generating correct workflow syntax from scratch”. This was our experience. Our rapid prototype of the RiboViz workflow in Nextflow, created as part of our comparative evaluation of our shortlisted systems, provided a sound basis for completing our implementation once we had decided upon Nextflow as our chosen workflow management system.

Rapid prototyping may not be suitable for the selection for all software or for all projects. For example, it would not be suitable for selecting software for large-scale IT projects or critical infrastructure. However, the use of rapid prototyping does offer a low-cost way of making a more informed selection of software to use within projects, than relying solely upon reviews and recommendations by others.

In conclusion, we agree that workflow management systems are a technology that “bioinformaticians need to be using right now” [3], and that they **can** implement right now using well-engineered open source tools.

## Supporting information

RiboViz workflow steps invoked when processing demultiplexed sample files

RiboViz workflow steps invoked when processing multiplexed sample files

## Acknowledgments

We thank our colleagues for discussions and comments on the manuscript: Hywel Dunn-Davies, Institute of Cell Biology, The University of Edinburgh; Alison Meynert, MRC Human Genetics Unit, The University of Edinburgh; Premal Shah, Department of Genetics, Rutgers University; Kit Transue, Seattle, USA. We thank the Software Sustainability institute for their support in starting this project.

## Supporting information

**S1 Fig. RiboViz workflow steps invoked when processing demultiplexed sample files.**

**S2 Fig. RiboViz workflow steps invoked when processing multiplexed sample files**.

