## Supplementary figures and images for "Using rapid prototyping to choose a bioinformatics workflow management system"

### RiboViz workflow steps invoked when processing demultiplexed sample files

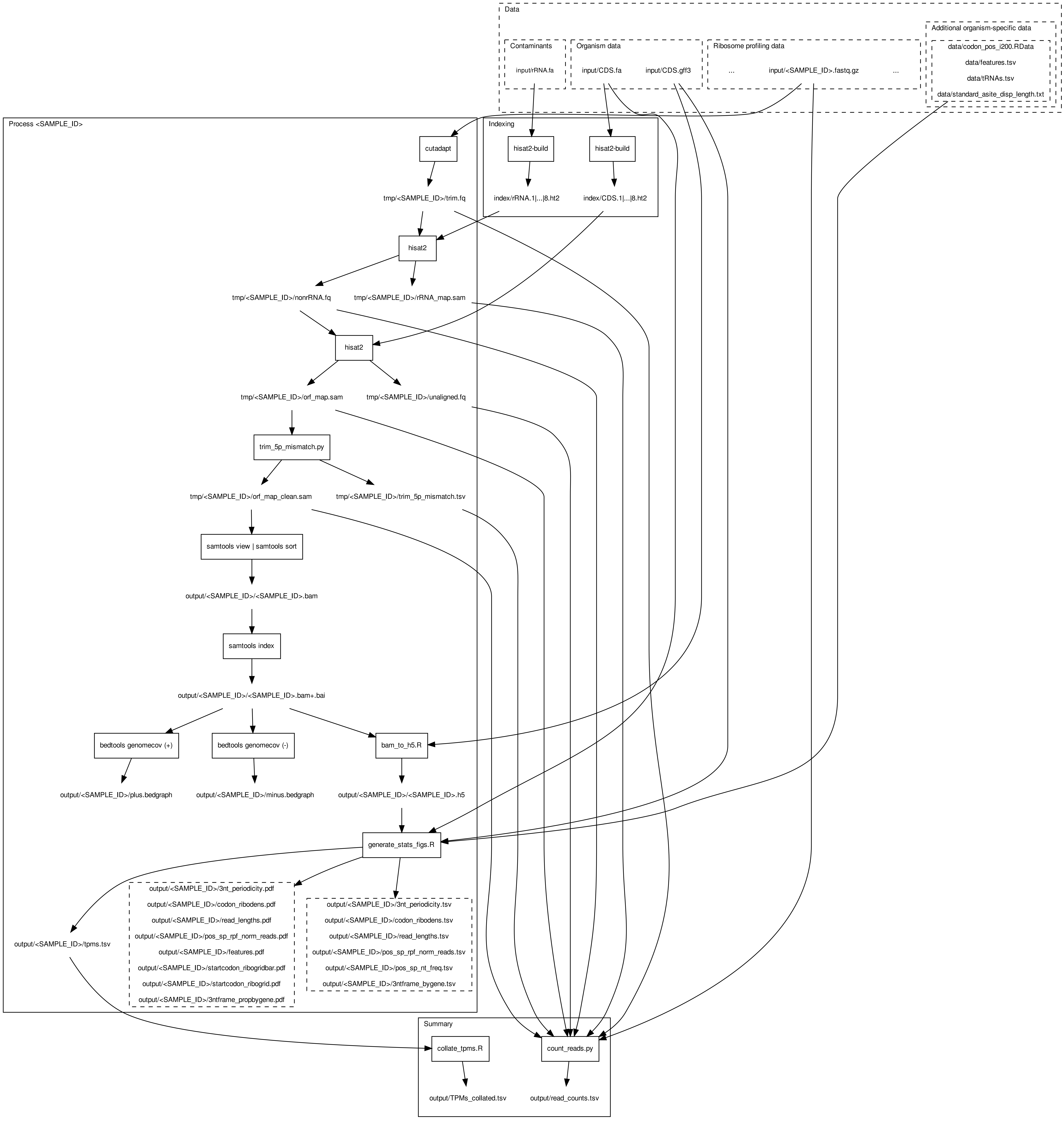

### RiboViz workflow steps invoked when processing multiplexed sample files

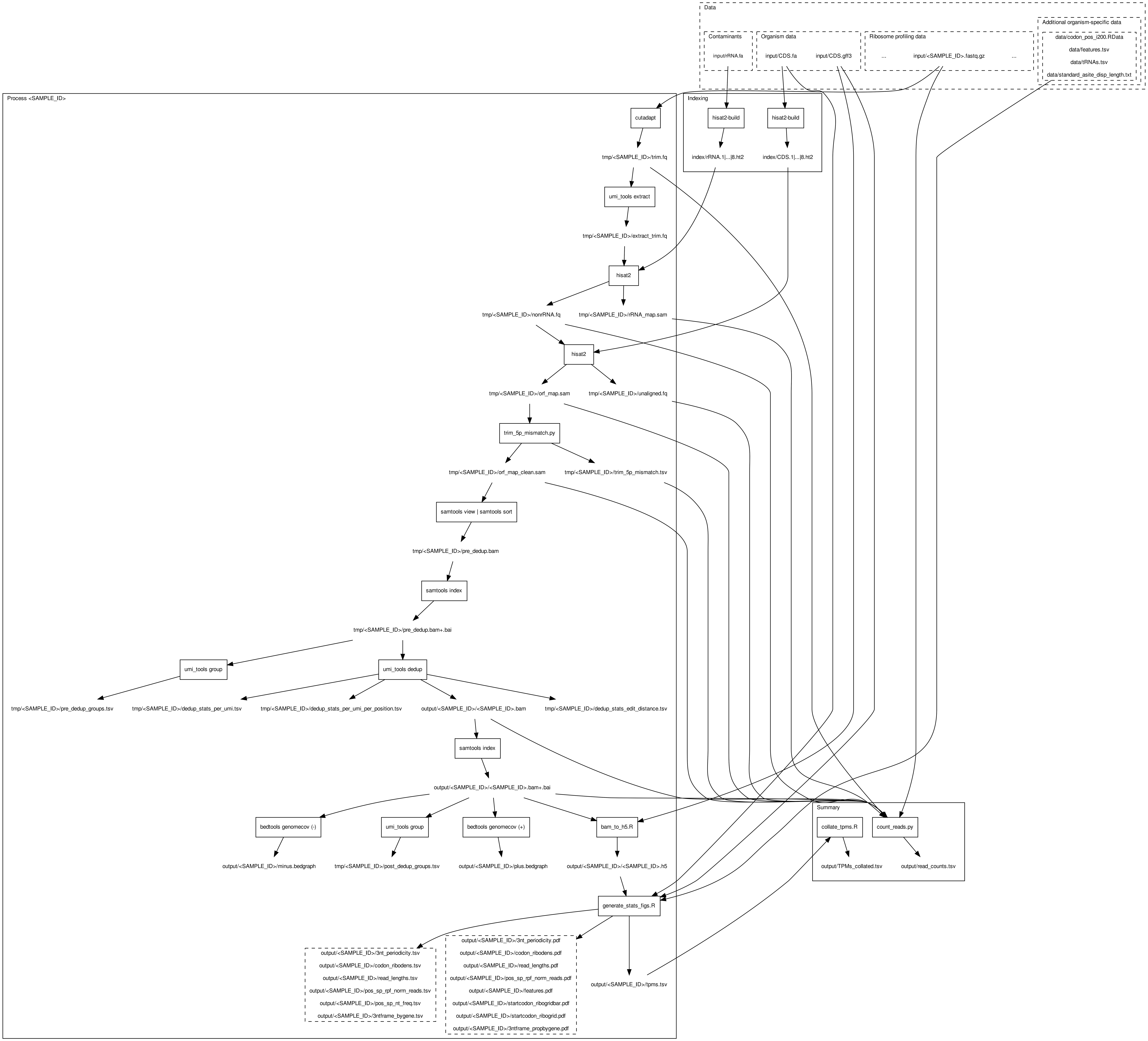
